# Genes with Specificity for Expression in the Round Cell Layer of the Growth Plate are Enriched in GWAS of Human Height

**DOI:** 10.1101/2021.02.24.432568

**Authors:** Nora E. Renthal, Priyanka Nakka, John M. Baronas, Henry M. Kronenberg, Joel N. Hirschhorn

## Abstract

Human adult height reflects the outcome of childhood skeletal growth. Growth plate (epiphyseal) chondrocytes are key determinants of height. As epiphyseal chondrocytes mature and proliferate, they pass through three developmental stages, which are organized into three distinct layers in the growth plate: 1) resting (round), 2) proliferative (flat), and 3) hypertrophic. Recent genome-wide association studies (GWAS) of human height identified numerous associated loci, which are enriched for genes expressed in growth plate chondrocytes. However, it remains unclear which specific genes expressed in which layers of the growth plate regulate skeletal growth and human height.

To connect the genetics of height and growth plate biology, we analyzed GWAS data through the lens of gene expression in the three dissected layers of murine newborn tibial growth plate. For each gene, we derived a specificity score for each growth plate layer and regressed these scores against gene-level p-values from recent height GWAS data. We found that specificity for expression in the round cell layer, which contains chondrocytes early in maturation, is significantly associated with height GWAS p-values (*p*=8.5×10^−9^); this association remains after conditioning on specificity for the other cell layers. The association also remains after conditioning on membership in an “OMIM gene set” (genes known to cause monogenic skeletal growth disorders, *p*<9.7×10^−6^). We replicated the association in RNA-seq data from maturing chondrocytes sampled at early and late time points during differentiation *in vitro*: we found that expression early in differentiation is significantly associated with *p*-values from height GWAS (*p*=6.1×10^−10^) and that this association remains after conditioning on expression at 10 days in culture and on the OMIM gene set (p<0.006). These findings newly implicate genes highlighted by GWAS of height and specifically expressed in the round cell layer as being potentially important regulators of skeletal biology.

## INTRODUCTION

GWAS of human height have identified numerous height-associated loci to date, making strides toward understanding the genetic underpinnings of human growth. As shown by our group and others, genes implicated by height GWAS are enriched for genes expressed in growth plate chondrocytes, the key cellular determinant of height [1]. However, the contribution of the specific stages of chondrogenic development have yet to be explored in relation to human height GWAS. Such an investigation offers the potential to advance our understanding of the developmental stages during growth plate maturation, since the comparison of growth plate zone gene expression and GWAS height associated genes could reveal important candidate genes and biological pathways missed in standard analysis of either methodology alone, and could also highlight a particular stage of chondrocyte maturation in the regulation of adult height.

Epiphyseal chondrocytes elongate bone by passing through three developmental stages in the growth plate: 1) resting (round), 2) proliferative (flat), and 3) hypertrophic. Regulation of each stage of chondrocyte development is paramount. Clues as to the mechanisms of regulation and the function of each cellular layer come from the specificity of gene expression within each zone. For example, many collagen family genes are highly expressed throughout all three layers of the growth plate, whereas matrix metalloproteinases, critical for the proteolytic processing of collagen and initiation of bone mineralization, are expressed preferentially in the hypertrophic zone.

In contrast to the proliferative and hypertrophic zones, the function of the resting zone has been historically less understood but has recently emerged as an area of active study [2-4]. Despite its “resting” nomenclature, this zone of the growth plate is now known to actively maintain the growth plate by supplying the cells that make up with proliferative zone [3]. These cells have been shown to express parathyroid hormone-related protein (PTHrP), which in turn, increases the activity of SOX9 and decreases the activity of Mef2c to maintain chondrocyte proliferation and delay hypertrophy in a non-cell autonomous manner [2, 5]. However, PTHrP+ cells are just one of several cell subgroups thought to be organized within the resting zone, concertedly contributing to growth plate renewal throughout childhood [2]. Thus, more studies regarding this critical layer are necessary, ideally including gene expression in the growth plate of the healthy child.

Of course, a natural challenge that currently precludes the ideal studies of human growth plate gene expression is the sparse availability of the large numbers of tissue samples from healthy pediatric growth plate that would be necessary to draw statistically powerful inferences. As an alternative approach, the current study harnesses the power of GWAS data from ∼700,000 individuals, in combination with gene expression data from dissected growth plate regions of healthy newborn mice and chondrocytes across different stages of development. Mouse models have long been used to study endochondral ossification. The value of such studies is emphasized by the conservation of phenotypes between human patients with monogenic disorders of skeletal growth and mice with knockouts of the orthologous genes, as well as the similarities between the morphologic changes during maturation of growth plate chondrocytes in the two species.

In the present study, we incorporate data from the largest published GWAS of height to date (N∼700,000) and two datasets relevant to growth plate gene expression: 1) newly published array data from microdissected layers of the murine growth plate [6] and 2) RNA sequencing of *in vitro* maturing chondrocytes. In contrast to our prior investigations, which focused on comparisons between growth plate expression and predefined lists of genes from a smaller GWAS [1], in the current investigation we consider the full set of results from a larger GWAS and data from all genes expressed in each layer of the growth plate. By using this approach, we are able to broadly measure the relationship between significance in height GWAS and expression patterns in the growth plate and show that genes highlighted by height GWAS are significantly enriched for genes expressed in the round layer and early in chondrocyte maturation. By integrating the GWAS and expression data, we can highlight genes and pathways expressed at early stages of chondrocyte maturation and associated with height in GWAS as being strong candidates to influence the genetic regulation of human height.

## RESULTS

### Genes expressed more specifically in the round layer of the growth plate are enriched for associations with height in GWAS

To understand which stages in chondrocyte development are relevant for regulation of human height, we integrated two different chondrocyte gene expression data sets with the results from GWAS of height: (1) microarray expression data from microdissected murine tibial growth plate (Figure 1A, [6]) and (2) RNAseq expression data from differentiated chondrocytes *in vitro* (Figure 1D, Supplemental Figure 1). Our first objective was to test for an association between gene expression in different dissected layers of the murine growth plate and genes highlighted by height GWAS. To generate a list of genes from GWAS, we used the software package MAGMA [7] to compute gene-level *p*-values using published GWAS summary statistics from a recent meta-analysis of height in ∼700,000 European individuals (6). To determine specificity of gene expression within each epiphyseal layer, we analyzed gene expression microarray data from murine growth plate microdissections, previously published by our research group. After normalization of gene expression levels (see Methods), specificity of gene expression within each epiphyseal layer was calculated as the percent of total expression attributable to each layer for each gene: specificity of 0% for a given gene within a layer indicates that none of the total expression of the gene can be attributed to that layer, whereas a specificity of 100% indicates the gene is exclusively expressed within that layer. As shown in Figure 1A, most genes are evenly expressed throughout all three layers of the growth plate, whereas others are more exclusively expressed in specific layers, likely related to the gene’s function in that growth plate zone. Taking the PTH signaling family as an example, we found that *Pthrp* is highly specific to the round zone, in which 72% of the growth plate’s *Pthrp* expression is found (vs. 18% flat and 10% hypertrophic). By contrast, the PTH receptor, *Pth1r*, is expressed predominantly in the hypertrophic zone (57%, vs. 8% round and 36% flat).

**Figure 1.**
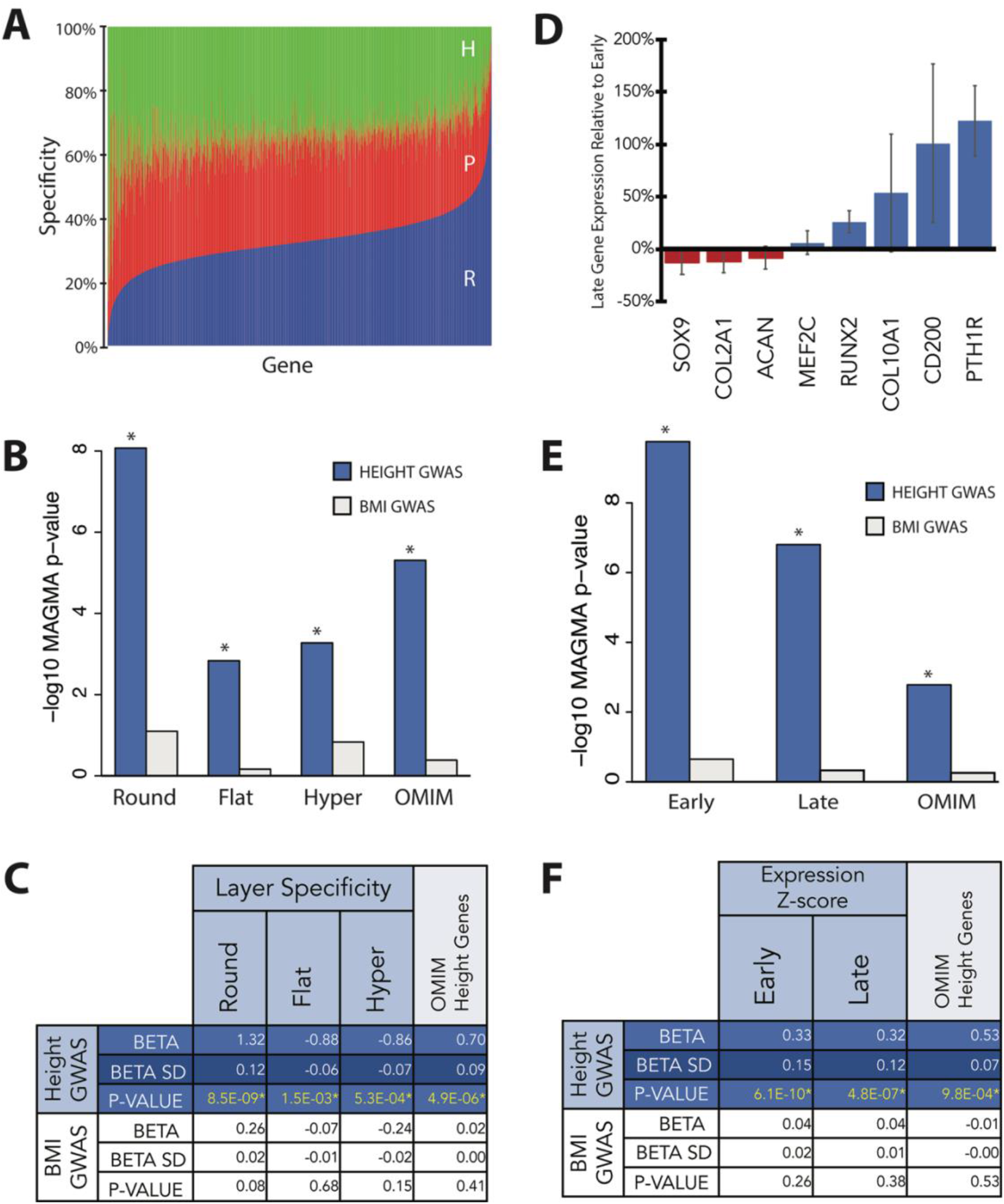
Gene expression in growth plate chondrocytes is associated with significance in GWAS for height. A) Specificity for each epiphyseal layer in the growth plate was calculated as the percent of total expression attributable to each layer for each gene. Round layer (blue), Proliferative (red), hypertrophic (green). B) Gene property association tests were performed between z-scores for GWAS-based gene-level p-values for height (dark blue) and BMI (grey) and 4 covariates: growth plate layer specificity and an OMIM gene set. C) Beta, standard deviation (SD), and p-values of the association between growth plate gene expression specificity and GWAS gene scores for height and BMI. Asterisks (*) indicate that a p-value passes Bonferroni correction for the number of tests (p <= 0.0015). D) Gene expression in differentiated chondrocytes in vitro, assayed by qPCR, normalized to 18s housekeeping gene and graphed relative to gene expression in early chondrocytes. The pattern of expression confirms upregulation of hypertrophic genes (Mef2c, Runx2, Col10a1, CD-200, Pth1r) and downregulation of typical early chondrogenesis genes (Sox9, Col2a1, Acan). E) Gene property association tests were performed between GWAS gene scores for height (dark blue) and BMI (grey) and 3 covariates: gene expression in early and late in vitro chondrocyte maturation and an OMIM gene set. F) Beta, standard deviation (SD), and p-values of the association between gene expression in early and late chondrocyte development in vitro and GWAS gene scores for height and BMI. Asterisks (*) indicate that a p-value passes Bonferroni correction for the number of tests (p <= 0.0017).

We then performed association tests between GWAS-based gene-level p-values (“GWAS gene scores”) and specificity scores for gene expression, using MAGMA to account for confounders such as gene length. Specificity scores for gene expression in each growth plate layer are strongly associated with height GWAS gene scores (all *p*<0.0015). As a negative control, we did not observe any associations with GWAS gene scores from a similarly sized study of body mass index (*p>*0.05) [8] (Figure 1B).

Because genes expressed more specifically in each of the three murine growth plate layers were enriched among GWAS height-associated genes, and because specificity scores in the three layers are correlated, we next sought to determine whether the associations were due specifically to expression in any particular layer of the epiphysis. To test for independent associations between height GWAS gene scores and gene expression in the growth plate, we conditioned on specificity for each layer of the growth plate. We found that conditioning on specificity for the round cell layer greatly attenuated the association between height GWAS gene scores and specificity scores for the other two layers (flat and hypertrophic, Figure 2A, Supplemental Table 1, all *p*>0.004, equivalent to *p*>0.05 after Bonferroni correction for 12 tests). However, the association with the round layer and height GWAS gene scores retained its significance even after conditioning on specificity for either of the other two layers (Figure 2A, Supplemental Table 1, all *p*<4.1×10^−6^). These observations indicate that the round cell layer is the only epiphyseal layer for which specificity scores are strongly and independently associated with height GWAS gene scores.

**Figure 2.**
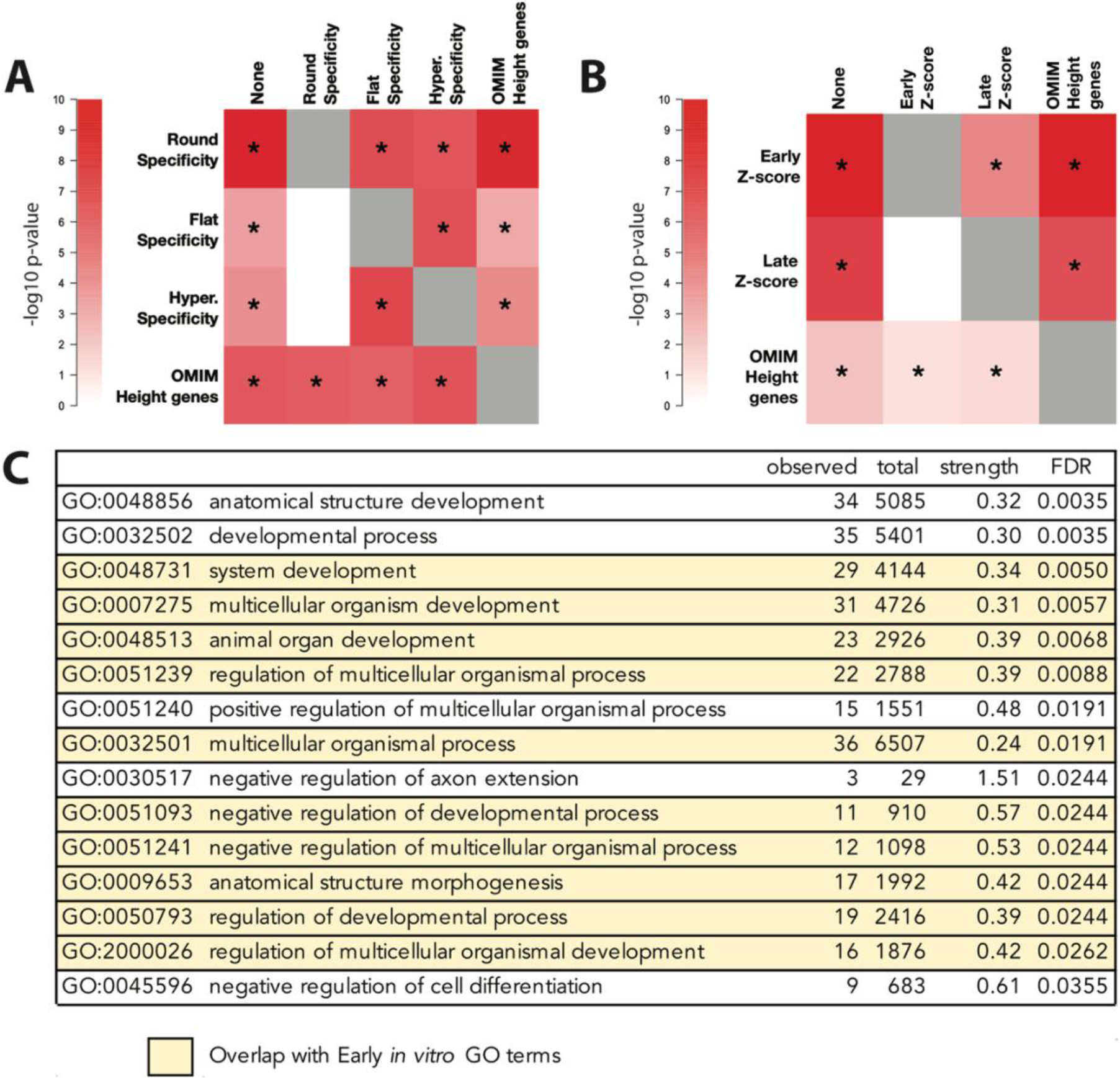
Round layer and early chondrocyte gene expression drive the association with height GWAS. A) To test for independent associations between height GWAS gene scores and gene expression in the growth plate, we conditioned on specificity for each layer of the growth plate and the OMIM height genes in turn. Conditioning on round cell layer specificity caused the association between height GWAS gene scores and specificity for the other two layers (flat and hypertrophic) to lose significance (p > 0.004). No other covariates demonstrated this pattern, suggesting that the round cell layer is the only epiphyseal layer independently associated with height GWAS gene scores. Conditioning on an OMIM height gene set did not decrease the round layer association. Asterisks (*) indicate that a p-value passes Bonferroni correction for the number of tests. B) We conditioned on expression at early and late chondrocyte developmental stages and an OMIM height gene set. Conditioning on early chondrocyte maturation gene expression caused the association between height GWAS gene scores and expression during late chondrocyte maturation to lose significance (p > 0.008). However, the association with early chondrocyte maturation expression does not lose significance when conditioning on late chondrocyte maturation expression, suggesting that gene expression in early chondrocyte development is independently associated with height GWAS gene scores. Conditioning an OMIM height gene set did not decrease the association. Asterisks (*) indicate that a p-value passes Bonferroni correction for the number of tests. C) 63 genes identified as having both high round layer specificity (normal transformed round layer specificity > 2 SD) and significance in height GWAS at or below the Bonferroni corrected p-value of 2.96×10^−6^ were tested for enrichment in known expression pathways and gene sets. Pathways that overlap with GO terms enriched in the 44 genes found to be highly expressed during early chondrocyte development (z-score Early expression > 2 SD) and significant in height GWAS at or below the Bonferroni corrected p-value of 2.96×10^−6^) are highlighted in yellow. The strength of pathway enrichment was calculated as Log_10_ (observed/expected) to determine the ratio between the number of genes annotated in our data compared to the expected count in a random network of the same size. Significance of pathway enrichment was estimated by calculating the False Discovery Rate (FDR). Shown are p-values corrected for multiple testing within each category using the Benjamini–Hochberg procedure.

### Known genes affecting skeletal growth are independently associated with height in GWAS

We also explored the relationship between specificity scores and membership in a set of 287 genes known to cause monogenic skeletal growth disorders (“OMIM genes”). Consistent with earlier studies [8], we find that the OMIM gene set is significantly associated with height GWAS gene scores (Figure 2A, Supplemental Table 1, *p* = 4.9×10^−6^). To test whether the association with OMIM genes is independent of the association with specificity scores, we conditioned on OMIM genes for round cell specificity and on round cell specificity for OMIM genes. We found that conditioning on the set of OMIM height genes did not attenuate the round layer association (Figure 2A, Supplemental Table 1, *p*=1.1×10^−8^) and conditioning on round layer specificity did not attenuate the OMIM gene set association (Figure 2A, Supplemental Table 1, *p*=6.9×10^−6^). We also find that specificity for the round cell layer does not differ significantly between the OMIM height genes and genes that are not in the OMIM list (Wilcoxon rank sum *p* = 0.5525; Supplemental Table 2). Thus, the association between round cell specificity and height gene scores is largely independent of the presence of known skeletal disease genes within the genes with higher round layer-specific expression.

### Validation of enrichment for early-expressed genes in GWAS height associations, using an *in vitro* model of chondrocyte differentiation

We next asked whether the association observed between murine growth plate gene expression and human height GWAS could be replicated in expression data from a different biologically relevant system. To assess the generalizability of our initial observations, we performed RNA-sequencing at different stages of maturation in a *v-myc* immortalized murine chondrogenic cell line (3, 4). Cells were cultured at high density in chondrogenic media for 3 days (early time-point) or 10 days (late time-point), prior to RNA extraction and sequencing. To confirm that these cells show the expected changes in gene expression during maturation, we performed qPCR for genes known to be expressed in differentiated chondrocytes. As seen in Figure 1D, the pattern of expression in late maturation confirms upregulation of hypertrophic genes (*Mef2c, Runx2, Col10a1, CD-200, Pth1r*) and downregulation of typical early chondrogenesis genes (*Sox9, Col2a1, Acan*), as compared to the pattern seen at the early time point. In the RNA-seq data, we observed 384 genes that were differentially expressed (>2 log fold change [LFC], FDR < 0.05) between early and late time points (Supplementary Figure 1). We observed significant associations between z-scores for gene expression at both timepoints of chondrocyte development and GWAS gene-level p-values for height (all *p*<4.8×10^−7^), but not for BMI (all *p*>0.05, Figure 1E & F). To test whether gene expression at a particular timepoint during *in vitro* maturation of chondrocytes is associated with height GWAS gene scores, we again performed conditional analyses and found that conditioning on early chondrocyte gene expression attenuated the association between height GWAS gene scores and expression during late chondrocyte maturation (Figure 2B, Supplemental Table 3, *p >* 0.008, equivalent to *p*>0.05 after Bonferroni correction for 6 tests). By contrast, the association with early chondrocyte maturation expression remained significant when conditioning on late chondrocyte maturation expression (*p*<3.1×10^−5^, Supplemental Table 3).

These observations are consistent with our earlier findings that the round cell layer was driving the association with height GWAS genes. The results thus further support the suggestion that gene expression in early chondrocyte development is responsible for the association with height GWAS gene scores. As was the case for the dissected growth plate gene expression data, conditioning on the OMIM gene-set also did not attenuate the association with expression at the early timepoint (*p*=3.4×10^−9^, Supplemental Table 3) and conditioning on the early timepoint did not attenuate the OMIM gene set association (*p*=0.006, Supplemental Table 3). We also found that OMIM height genes have significantly higher expression during the early timepoint than non-OMIM genes (Wilcoxon rank sum *p* = 2.34 × 10^−7^; Supplemental Table 2). Thus, though known height genes are enriched for genes that are highly expressed during early maturation, our conditional analysis shows the association between early expression and height GWAS gene scores is not completely driven by genes known to underlie monogenic skeletal growth disorders.

### Pathway analysis reveals enrichment in processes governing chondrocyte maturation and development

Within the round cell layer, specific expression of sets of related genes (pathways) likely promotes the maintenance of the growth plate and influence subsequent skeletal growth. We therefore asked whether specific genes or pathways expressed within the round cell layer underpin the association observed with height GWAS. Sixty-three genes were identified as having both high round layer specificity (normal transformed round layer specificity > 2 SD) and gene-level *p* values in height GWAS at or below the Bonferroni corrected significance threshold of 2.96×10^−6^ (Supplemental Table 4).

These genes were tested for enrichment in known expression pathways and gene sets (Figure 2C). Fifteen gene ontology terms for biological processes showed a strong enrichment in pathways known to govern cell development, including members of the Wnt signaling (NFATC2, FOSL1, DKK3, FOSL1, NFATC2, TNS2, and WNT9A), parathyroid hormone signaling (PTHLH, also known as PTHrP) pathways, and modifiers of the TGFbeta-superfamily (LTBP4 and THSD4). Only two were also found within our curated OMIM height gene list: parathyroid hormone like hormone (PTHLH, human 12p11.22, OMIM *168470, known to cause Brachydactyly, type E2 [9]) and the member of the RAS oncogene family 23 (RAB23, human 6p12.1-p11.2, OMIM *606144, known to cause Carpenter syndrome [10]). Other implicated genes include those involved in the formation of growth plate cartilage extracellular matrix, including structural proteins in the cartilage matrix (COL14A1, COL5A3), as well as extracellular matrix protein 1 (ECM1) Involved in endochondral bone formation as a negative regulator of bone mineralization. Other genes among the 63 identified have not yet been associated with known regulatory pathways in growth plate biology (Supplemental dataset 1) and thus represent novel candidate genes, potentially directing the growth of bones in mouse and human.

Interestingly, when we performed a similar analysis of the genes driving the association between early chondrogenesis and height GWAS, we identified 44 genes highly expressed at Day 3 in cell culture (gene expression z-score > 2 SD) and significant in height GWAS at or below the Bonferroni corrected p-value of 2.96×10^−6^. These genes were likewise tested for enrichment in known expression pathways. Although no genes directly overlapped between the 63 (GWAS + Round) and 44 (GWAS + Early) sets, there was significant overlap of the enriched GO pathways governing organismal development and regulation in both groups. Ten of the fifteen (10/15) gene ontology terms found to be enriched in the genes driving the association between the round layer and GWAS were also found to be enriched in the genes associating early chondrocyte development in vitro and GWAS (Figure 2C, highlighted in yellow). Additionally, of the 63 genes identified as having both high round layer specificity and significance in height GWAS, 20 were found to be differentially expressed between early and late timepoints of maturation to hypertrophy *in vitro*, further suggesting that these genes may play a role in the maturation of the growth plate chondrocyte (Supplemental Table 5).

## DISCUSSION

This study demonstrates a strong association between growth plate gene expression patterns and genes implicated by GWAS of human height. Furthermore, our results suggest that gene expression in the round cell layer is particularly important to prioritizing and validating genes from human height GWAS, and implicate gene expression within the round layer/early chondrocyte maturation as a key determinant of skeletal growth. Pathways underpinning this connection underscore the importance of known developmental gene networks for chondrocyte development (such as PTHrP, BMP, and WNT signaling), as well as gene expression within the round cell layer that directs cell-cell interactions and cell structure. By intersecting genes expressed early in maturation with height GWAS genes, we have highlighted new genes as candidates for regulating human skeletal growth through actions in the growth plate.

Our studies illustrate how GWAS gene prioritization can be augmented by evaluating gene expression within a tissue of biological relevance, especially if data from relevant tissues are available. In the present study, we focused on analysis of GWAS with expression data generated by our laboratories; however, other data sets could be used for future analyses of this type. As a proof of principle, we performed similar analysis of publicly available microarray data from dissected embryonic murine tibiae, partitioned into three general regions [11]. Although these partitioned regions do not directly parallel the zones of the post-natal growth plate, we were encouraged to find a trend congruent with the findings described here: regions representing earliest chondrocyte development (termed 1 and 2 by James et al.) are significantly associated with height and remain significant after conditional analysis (Supplemental Table 6). Recently developed methods to prioritize genes from GWAS results offer the possibility to include many data sets simultaneously and identify those which are most helpful with gene prioritization [12]. Thus, the methodology described here provides an important technical resource for comparison of gene expression in different stages of chondrocyte development and will aid in the ongoing translation of GWAS associated SNPs to implicated causal genes for regulating skeletal growth.

The current study represents an advance over prior investigations, with larger GWAS datasets and updated gene expression data from tissues of more specific biological relevance: growth plate zones and maturing chondrocytes. Additionally in this study, we utilized the full set of expression data and GWAS summary association statistics in our analyses. We found significant, independent associations between height GWAS and the round cell layer and early timepoints in chondrocyte maturation, further identifying 63 genes acting in the round cell layer to influence the genetic regulation of height. Among these, reassuringly, were *Pthrp*, members of the WNT signaling pathway, and collagen family genes. Potentially interesting targets, less studied in the growth plate, were genes such as neuroblastoma suppressor of tumorigenicity 1 (*Nbl1*), the founding member of the DAN family of transforming growth factor beta antagonists, shown to specifically antagonize BMP2 in the ovary [13] and found to be differentially expressed in cartilage from human osteoarthritis [14]. Likewise, Netrin-1 (*Ntn1*) is a secreted molecule of the laminin superfamily, best known for its role in axon guidance [15, 16], but it has also been shown to play a role in chondrocyte migration [17]. Further studies will be required to define the role of *Nbl1, Ntn1*, and other genes in the round zone. Nevertheless, they exemplify the quite interesting candidate genes that have emerged from our analysis and illustrate the rich potential of this methodology to narrow the list of GWAS genes for future functional studies.

Progressing from associations in GWAS studies to a set of causal genes remains a significant obstacle to fully harnessing the power of GWAS to reveal causal disease biology, even for a phenotype such as height where many relevant genes and pathways are known. One key step in this path is the identification of the cell type(s) and stage(s) of development in which causal genes exert their effects. The current study combines data from the largest published height GWAS to date and datasets of growth plate gene expression to identify the round zone as critical for human height determination. Continued improvement of methods to prioritize genes from GWAS, further increases in GWAS sample sizes, as well as improved methodologies to investigate gene expression or chromatin accessibility with greater tissue specificity (such as single cell methods) will all improve future efforts to identify genes that regulate skeletal growth. Our current studies contribute to this work by shedding new light on the biology of endochondral bone growth, highlighting potential new regulators of human height, and making new gene expression data for chondrocyte maturation available for future studies.

## MATERIALS AND METHODS

### Cell Culture

The v-myc immortalized chondrogenic murine cell line derived from <u>m</u>urine embryonic limb <u>b</u>ud at gestational day <u>13</u> (MLB13) (3, 4), was maintained in DMEM/F12 (Gibco, Grand Island, NY, USA) with 10% FBS (Gibco), 100 U/mL penicillin, and 100 mg/mL streptomycin sulfate (Life Technologies, Inc., Grand Island, NY, USA), at 37°C with 5% CO2. Cells were cultured at high density with the addition of 50 ug/mL ascorbate (Sigma-Aldrich) and 1x insulin-transferrin-selenium (ThermoFisher) for 3 days (early time-point) or 10 days (late time-point). Cells were harvested and RNA extracted for sequencing.

### RNA Extraction and Sequencing

RNA was extracted from our cultured chondrocyte cell line (four samples per time point) and purified on RNeasy microcolumns (Qiagen) per the manufacturer’s instructions. RNA with an A260/280 ratio less than 1.7 was not used for downstream analysis. RNA-sequencing (RNA-seq) was performed with rRNA-depletion via the KAPA HyperPrep RNA & Ribo-Erase kit by the Biopolymers Facility, Department of Genetics, Harvard Medical School. Briefly, rRNA was hybridized to probes and digested with enzymes. Remaining RNA was converted into cDNA, prior to library preparation and post-PCR Cleanup.

### Quantitative PCR

For cDNA synthesis, 1 μg RNA was used in synthesis reactions according to the instructions of the manufacturer (Primescript RT). SYBR Green-based quantitative PCR detection was performed using FastStart Universal SYBR Green (Roche) on a StepOne Plus (Applied Biosystems) thermocycler.

Transcript levels for genes of interest were determined relative to 18s housekeeping gene using standard methods [18]. All PCR primer sequences are listed in Table 1.

**Table 1.**
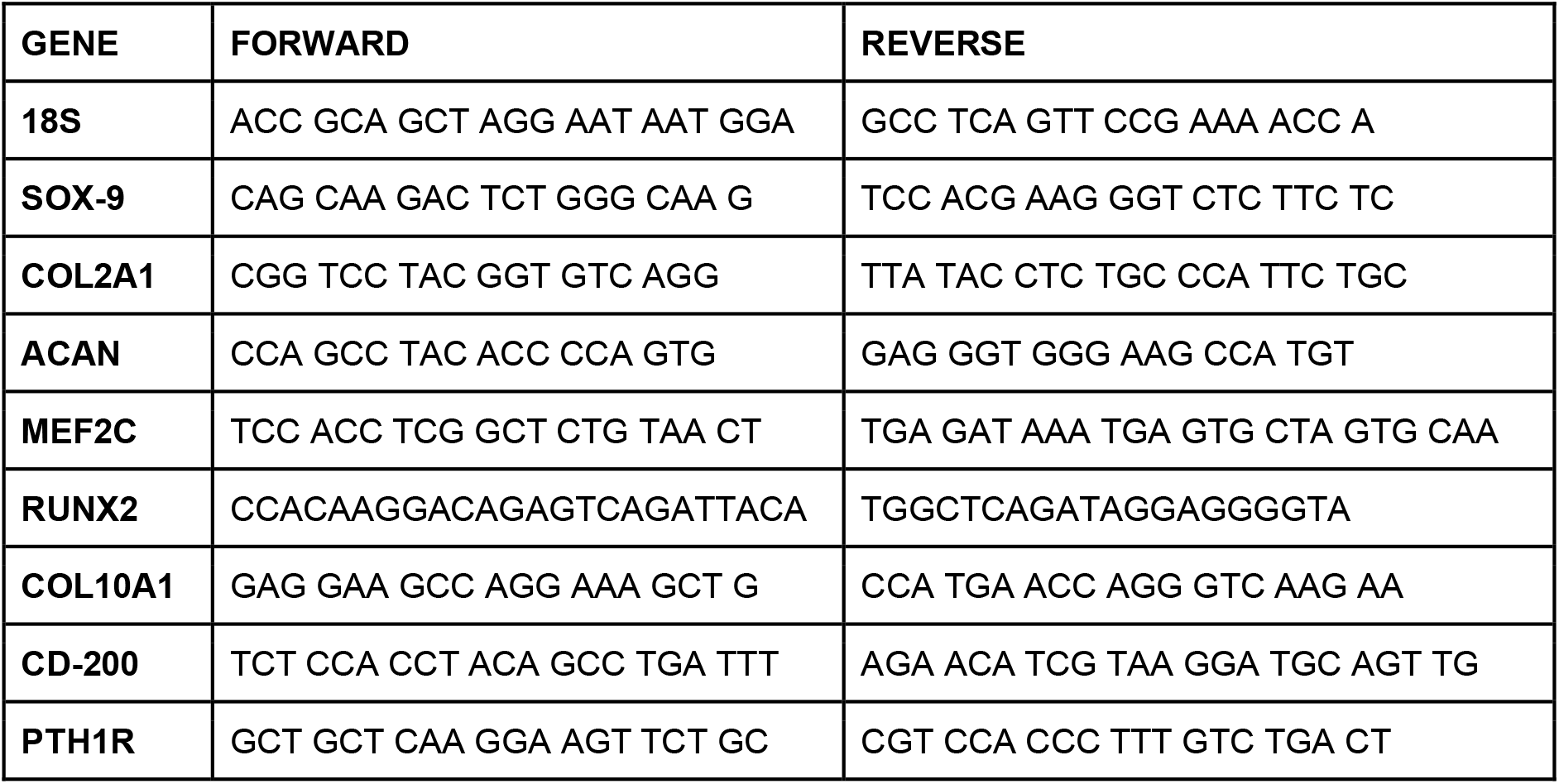
Primer sequences used for quantitative PCR.

### Mouse-to-human gene mapping

We downloaded published microarray data from growth plate dissections from the GEO data repository (https://www.ncbi.nlm.nih.gov/geo/; accession number: GSE87605). We mapped Affymetrix 430 2.0 probe identifications to mouse genes using the MGI database (Mouse Genome Informatics, The Jackson Laboratory; URL: http://www.informatics.jax.org/). We then mapped mouse gene names to their human homologs using the Ensembl database, accessed through the R (version 3.5) package BioMaRt (version 2.36.1) [19, 20].

### Calculation of epiphyseal layer expression specificity

Specificity for each epiphyseal layer was calculated as the proportion of expression for each gene; a specificity of 0 for a given gene and layer indicates that none of the total expression of the gene can be attributed to the layer and a specificity of 1 indicates that all expression of the gene can be attributed to the layer.

### Calculation of expression z-scores for chondrocyte maturation stage

We first obtained GC-content for each gene and normalized for GC content across all samples and time points of the RNA-seq dataset using the R package EDASeq (version 2.16) [21]. Gene expression was averaged across four samples for each time point: (1) day 3 or “early maturation stage” and (2) day 10 or “late maturation stage”. We then transformed gene expression to z-scores within each stage of chondrocyte maturation.

### Height GWAS p-values

We obtained SNP-level GWAS summary statistics from a recent large GWAS of height. https://portals.broadinstitute.org/collaboration/giant/index.php/GIANT_consortium_data_files. These data have been previously described [8]; briefly, the authors conducted a meta-analysis of height in ∼700,000 European individuals and identified 3290 genome-wide significant SNP loci associated with height. We analyzed 2,315,711 SNPs from this GWAS dataset mapping to 16,877 genes.

### Gene-set analysis using MAGMA

We used MAGMA (version 1.07) to evaluate association of gene-level height GWAS association statistics with epiphyseal layer specificities and time point specific z-scores in order to determine which layer or time-point is most associated with genes identified in height GWAS. We first computed gene-level p-values for height GWAS p-values [8] using MAGMA’s snp-wise mean model with 1000 Genomes European data for account for linkage disequilibrium (LD).

The layer and time-point specific metrics were calculated as described above (see “Calculation of epiphyseal layer expression specificity and z-scores” section). These metrics were then used as covariates in MAGMA’s gene property analysis method, which is a two-sided association test between covariates and the phenotype, height. For the microarray and RNA-seq datasets, we included as a covariate, membership in a gene set of Mendelian genes involved in skeletal development (obtained from OMIM), in order to account for the possibility that association with an epiphyseal layer can be explained by all expressed genes being a part of this gene set. By default, MAGMA also conditions on the gene size, gene density, sample size, and the inverse of the mean minor allele count in the gene and the log of these variables for every gene-set analysis.

### Conditional gene-set analysis using MAGMA

If two covariates (or gene properties) are significant in the basic gene property association analysis, the effect of one covariate may be largely due to the other covariate. In this case, the significance of one covariate will disappear after conditioning on the second. To detect these confounding variables, we performed conditional analysis on both datasets using MAGMA. For the microarray datasets, we conditioned on each of the three layers and the gene set of skeletal development genes separately. For the RNA-seq dataset, we conditioned on both time points as well as the gene set of skeletal development genes separately.

### Collation of an OMIM gene-set for Skeletal Disease

We curated a list of 287 genes (OMIM height gene set, Supplementary Data) in which mutations are known to cause a skeletal phenotype in humans by combining data from the Online Mendelian Inheritance in Man (OMIM) database and additional genes reported in the literature to cause human skeletal dysplasia. To include genes in which mutations cause abnormal skeletal growth due to local skeletal dysfunction, genes that impair growth through a systemic mechanism (such as growth hormone deficiency) were excluded.

### Pathway analysis of genes driving the association between the round layer, early chondrogenesis, and height GWAS

To identify genes and biological pathways driving the association between the round layer and height GWAS, we identified 63 genes as having both high round layer specificity (normal transformed round layer specificity > 2 SD) and significance in height GWAS at or below the Bonferroni corrected p-value of 2.96×10^−6^. Likewise, to identify genes and biological pathways driving the association between early chondrogenesis and height GWAS, we identified 44 genes as being highly expressed at Day 3 in cell culture (gene expression z-score > 2 SD) and significance in height GWAS at or below the Bonferroni corrected p-value of 2.96×10^−6^. These gene sets were tested for enrichment in known expression pathways. The strength of pathway enrichment was calculated as Log10 (observed/expected) to determine the ratio between the number of genes annotated in our data compared to the expected count in a random network of the same size. Significance of pathway enrichment was estimated by calculating the False Discovery Rate (FDR).

## ACKNOWLEDGMENTS

We would like to thank Dr. Shigeki Nishimori for the use of his published growth plate gene expression data set. This work was supported by the Eunice Kennedy Shriver National Institute of Child Health and Human Development (NICHD), the US National Institutes of Health (NIH), and Boston Children’s Hospital/Broad Institute Collaborative Project funding from the Boston Investment Conference.

## Data availability statement

The murine growth plate data that support the findings of this study are available in the GEO data repository (https://www.ncbi.nlm.nih.gov/geo/; accession number: GSE87605). These data were derived from the following resources available in the public domain: MGI database (Mouse Genome Informatics, The Jackson Laboratory; URL: http://www.informatics.jax.org/).

The GWAS summary statistics are available at https://portals.broadinstitute.org/collaboration/giant/index.php/GIANT_consortium_data_files.

The chondrocyte RNAseq data will be made openly available in the GEO data repository prior to publication.

---

**Supplemental Figure 1.**
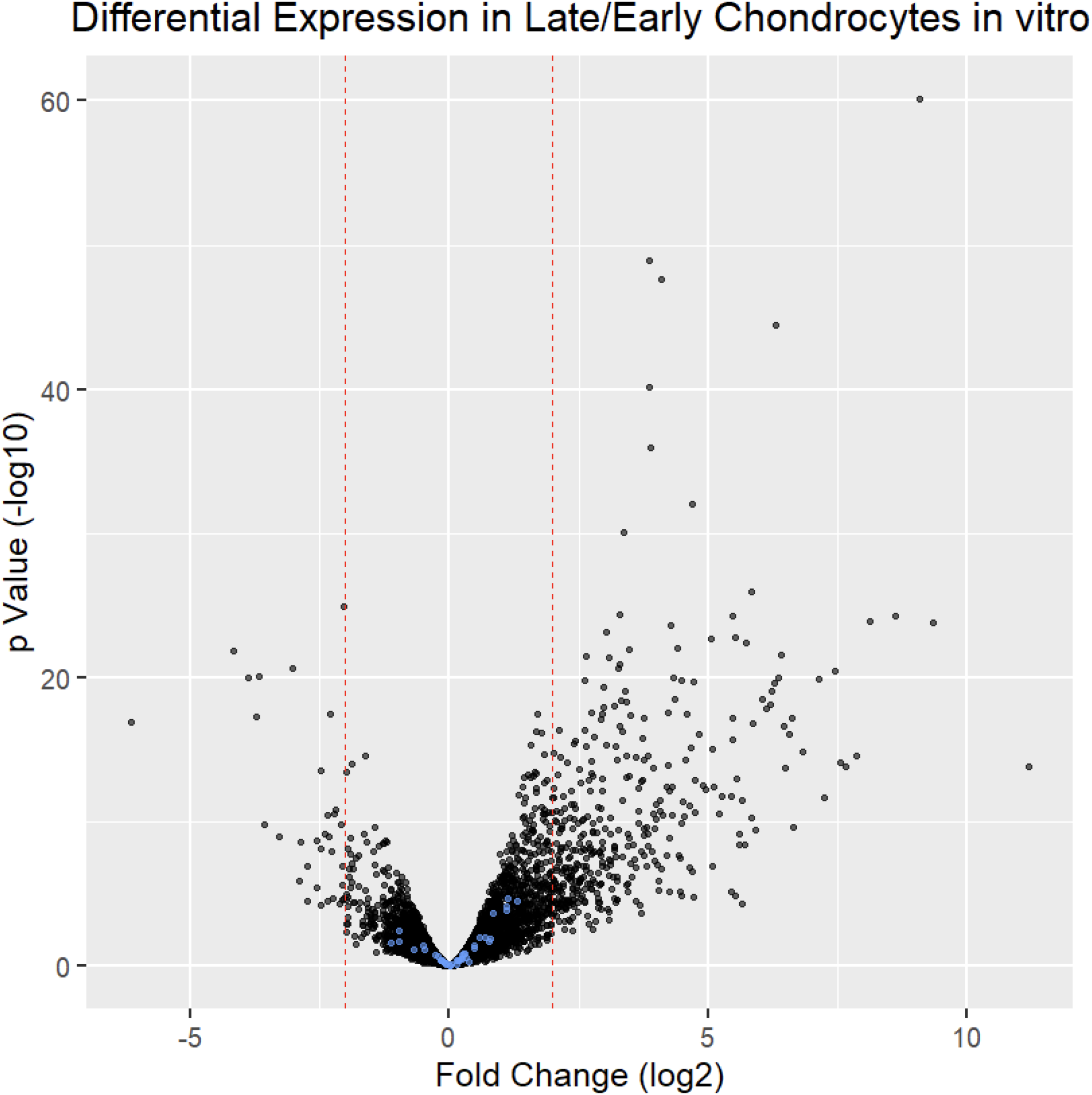
Immortalized chondrocytes were cultured at high density in chondrogenic media for 3 days (early time-point) or 10 days (late time-point), followed by RNA extraction and sequencing. Differential expression was analyzed with R and the Bioconductor R package DESeq. The 44 genes found to be highly expressed during early chondrogenesis (z-score Early expression > 2 SD) and significant in height GWAS at or below the Bonferroni corrected p-value of 2.96×10^−6^) are blue. Red dashed lines mark LFC +/-2.

**Supplemental Table 1.** This table shows the MAGMA association *p*-values used to generate Figure 2A. To test for independent associations between height GWAS gene scores and gene expression in the growth plate, we conditioned on specificity for each layer of the growth plate and the OMIM height genes in turn. We found that conditioning on specificity for the round cell layer caused the association between height GWAS gene scores and specificity for the other two layers (flat and hypertrophic) to lose significance (*p* > 0.004). No other covariates demonstrated this pattern, suggesting that the round cell layer is the only epiphyseal layer independently associated with height GWAS gene scores. Notably, conditioning on our set of OMIM height genes did not decrease the round layer association; therefore, the association is not due to the presence of known skeletal disease genes within our round layer specificity data set. Asterisks (*) indicate that a p-value passes Bonferroni correction for the number of tests.

| Conditioned Variable: | None | Round Layer | Flat Layer | Hypertrophic Layer | OMIM Genes |
| --- | --- | --- | --- | --- | --- |
| Round Layer | 8.47e-09* | NA | 1.49e-06* | 4.12e-06* | 1.18e-08* |
| Flat Layer | 1.46e-03* | 0.72 | NA | 2.29e-06* | 2.97e-03* |
| Hypertrophic Layer | 5.32e-04* | 0.83 | 8.65e-07* | NA | 3.34e-04* |
| OMIM Gene Set | 4.93e-06* | 6.91e-06* | 9.74e-06* | 3.12e-06* | NA |

**Supplemental Table 2.** To test whether known skeletal development genes from OMIM (“OMIM genes”) are over-represented in any growth plate layer or developmental timepoints, we performed two-sample Wilcoxon rank sum tests on specificity values and expression z-scores for OMIM genes and non-OMIM genes. We found that the OMIM genes had significantly higher expression z-scores during early and late development than non-OMIM genes. We found that OMIM genes had significantly lower specificity to the flat layer than non-OMIM genes. We also found that specificity to the round and hypertrophic cell layers does not differ significantly between OMIM genes and non-OMIM genes. Asterisks (*) indicate significance after Bonferroni correction for 5 tests (*p* < 0.01).

|  | Mean Specificity/Z-scores for OMIM Height Genes | Mean Specificity/Z-scores for non-OMIM Height Genes | Wilcoxon Rank Sum P-value for OMIM genes vs non-OMIM genes |
| --- | --- | --- | --- |
| Round Layer Specificity | 0.335 | 0.330 | 0.552 |
| Flat Layer Specificity | 0.305 | 0.331 | $2.277 \times 10^{-6*}$ |
| Hypertrophic Layer Specificity | 0.359 | 0.338 | 0.192 |
| Early Timepoint Expression Z-Score | 0.716 | -0.014 | $2.340 \times 10^{-7*}$ |
| Late Timepoint Expression Z-score | 0.683 | -0.014 | $7.383 \times 10^{-11*}$ |

**Supplemental Table 3.** This table shows the MAGMA association p-values used to generate Figure 2B. To test for independent associations between height GWAS gene scores and gene expression in the growth plate, we conditioned on expression at early and late chondrocyte developmental stages and the OMIM height genes in turn. We found that conditioning on early chondrocyte maturation gene expression caused the association between height GWAS gene scores and expression during late chondrocyte maturation to lose significance (*p* > 0.008). However, the association with early chondrocyte maturation expression does not lose significance when conditioning on late chondrocyte maturation expression, suggesting that gene expression in early chondrocyte development is independently associated with height GWAS gene scores. Again, conditioning an OMIM height genes set did not decrease the association, suggesting the association is not due to the presence of known skeletal disease genes during early maturation. Asterisks (*) indicate that a p-value passes Bonferroni correction for the number of tests.

| Conditioned Variable: | None | Early Timepoint | Late Timepoint | OMIM Genes |
| --- | --- | --- | --- | --- |
| Early Timepoint | 6.14e-10* | NA | 3.13e-05* | 3.35e-09* |
| Late Timepoint | 4.78e-07* | 0.036 | NA | 2.04e-06* |
| OMIM Gene Set | 9.78e-04* | 0.006* | 4.55e-03* | NA |

**Supplemental Table 4:**
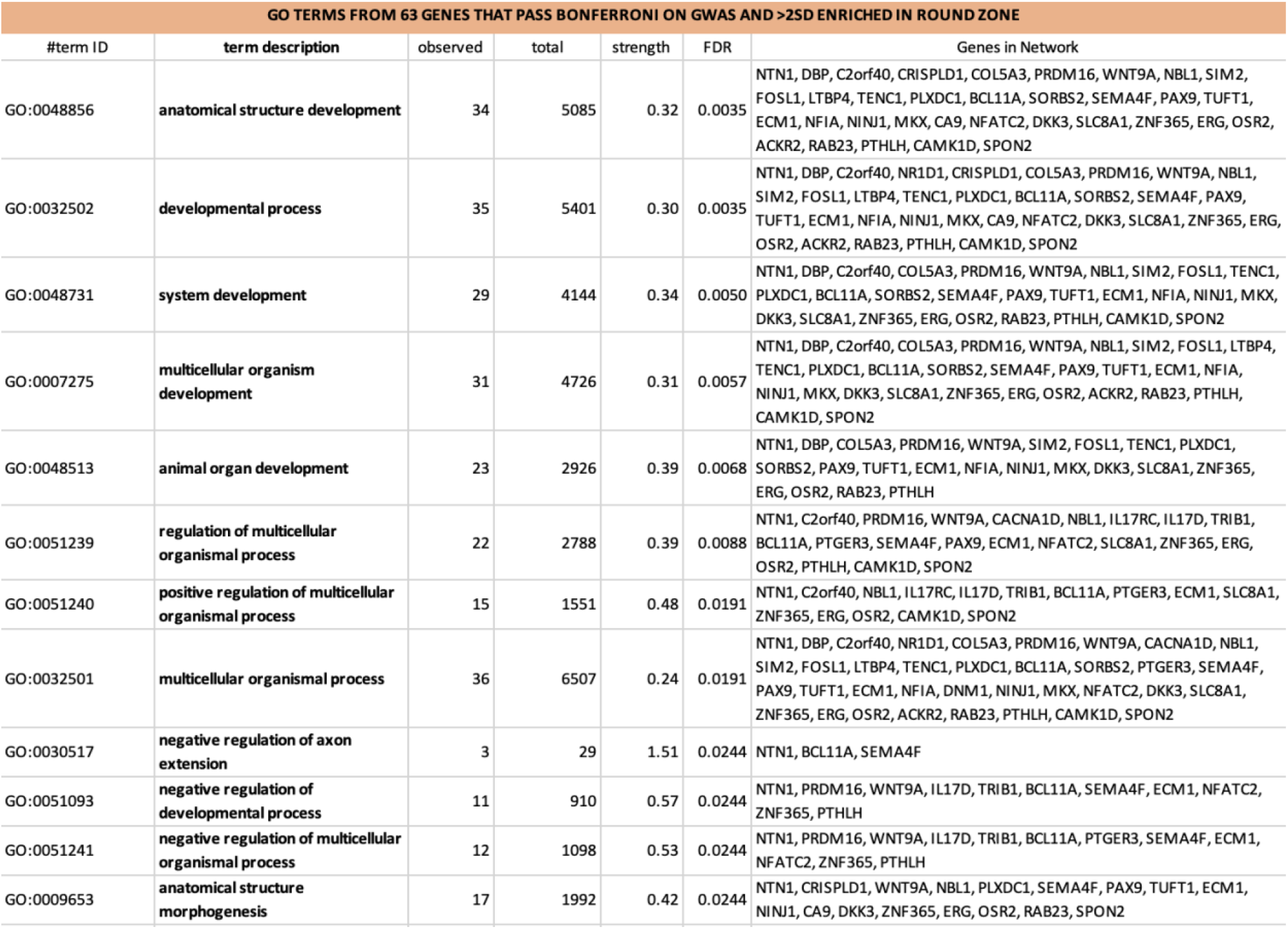
Full gene list information for the Gene ontology terms of the 63 genes identified as having both high round layer specificity (normal transformed round layer specificity > 2 SD) and significance in height GWAS at or below the Bonferroni corrected p-value of 2.96×10^−6^ were tested for enrichment in known expression pathways and gene sets.

**Supplemental Table 5:**
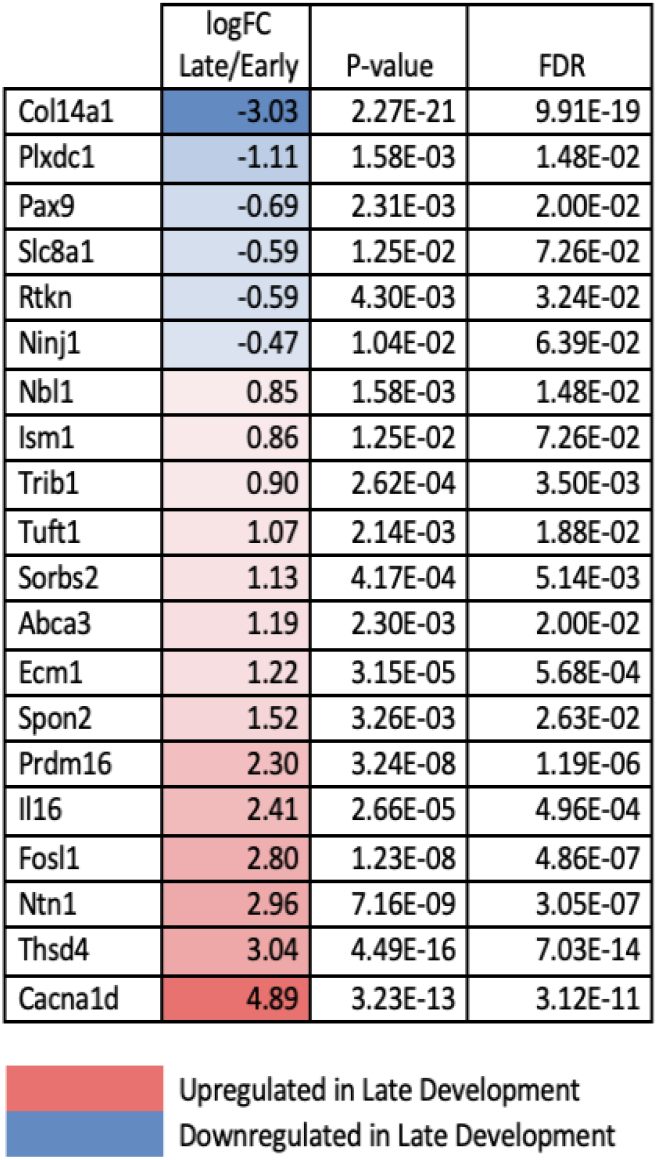
Of the 63 genes identified as having both high round layer specificity (normal transformed round layer specificity > 2 SD) and significance in height GWAS at or below the Bonferroni corrected p-value of 2.96×10^−6^, 50 were expressed by our chondrocyte cell line and 20 were found to be regulated during maturation to hypertrophy *in vitro*.

**Supplemental Table 6:** (A) We find significant associations between specificity for zones 1 and 2 (*p* < 0.01) and height GWAS gene scores. Beta, standard deviation (SD), and p-values of the association between growth plate zone expression specificity from a published dataset [11] and GWAS gene scores for height. (B) To test for independent associations between height GWAS gene scores and gene expression in the growth plate, we conditioned on specificity for each zone of the growth plate. We find that Zones 1 and 2 both remain significantly associated with height GWAS even after conditioning on the other two zones. Asterisks (*) indicate that a p-value passes Bonferroni correction for the number of tests.

|  | Height association p-values |  |  |
| --- | --- | --- | --- |
| Variable | BETA | BETA SD | p-value |
| Zone 1 | 4.76E+00 | 5.48E-02 | 6.28E-03* |
| Zone 2 | -4.66E+00 | -5.70E-02 | 4.60E-03* |
| Zone 3 | 2.76E-01 | 4.09E-03 | 8.38E-01 |
| OMIM Gene Set | 1.13E+00 | 9.23E-02 | 2.65E-06* |

| Conditioned Variable | Zone 1 | Zone 2 | Zone 3 |
| --- | --- | --- | --- |
| Zone 1 | NA | 2.76E-02 | 3.24E-04* |
| Zone 2 | 2.00E-02 | NA | 4.03E-04* |
| Zone 3 | 1.90E-02 | 3.33E-02 | NA |

## Notes

### Competing Interest Statement

The authors have declared no competing interest.

